# The *Drosophila* ERG channel *seizure* plays a role in the neuronal homeostatic stress response

**DOI:** 10.1101/521005

**Authors:** Alexis S. Hill, Poorva Jain, Yehuda Ben-Shahar

**Affiliations:** Department of Biology, Washington University in St. Louis, St. Louis, Missouri, United States of America; Deparment of Biology, College of the Holy Cross, Worcester, Massachusetts, United States of America

**Author notes:** Corresponding Author (YB-S).

## Abstract

Neuronal physiology is particularly sensitive to acute stressors that affect excitability, many of which can trigger seizures and epilepsies. Although intrinsic neuronal homeostasis plays an important role in maintaining overall nervous system robustness and its resistance to stressors, the specific genetic and molecular mechanisms that underlie these processes are not well understood. Here we used a reverse genetic approach in *Drosophila* to test the hypothesis that specific voltage-gated ion channels contribute to neuronal homeostasis, robustness, and stress resistance. We found that the activity of the voltage-gated potassium channel *seizure (sei)*, an ortholog of the mammalian ERG channel family, is essential for protecting flies from acute heat-induced seizures. Although *sei* is broadly expressed in the nervous system, our data indicate that its impact on the organismal robustness to acute environmental stress is primarily mediated via its action in excitatory neurons, the octopaminergic system, as well as glia. Furthermore, our studies suggest that human mutations in the human ERG channel (hERG), which have been primarily implicated in the cardiac Long QT Syndrome (LQTS), may also contribute to the high incidence of seizures in LQTS patients via a cardiovascular-independent neurogenic pathway.

**Author Summary:** Neurons are extremely sensitive to diverse environmental stressors, including rapid changes in the ambient temperature. To buffer stress, all animals have evolved diverse physiological mechanisms to protect neuronal activity from acute and chronic stressors, and failures of these safeguards often lead to hyperexcitability, episodic seizures, and chronic epilepsy. Although seizures and related syndromes are common, their underlying molecular and genetic factors, and their interactions with environmental triggers, remain mostly unknown. Here, we show that in the fruit fly, mutations in the ERG voltage-gated potassium channel *seizure (sei)*, an ortholog of the human hERG channel that has been previously implicated in the cardiac Long-QT syndrome, could also increase seizure susceptibility. We demonstrate that in addition to its cardiac expression, the *sei* channel is broadly expressed in the nervous system, specifically localized to axonal projections, and is specifically required in excitatory and modulatory neurons, as well as non-neuronal glia for maintaining organismal resistance to heat-induced seizures. Thus, our work suggests that the previously reported increase in seizure susceptibility in individuals with mutations in hERG are likely directly related to its neuronal action, independent of its cardiac function.

## Introduction

Neuronal homeostatic responses to acute and long-term environmental stressors are essential for maintaining robust behavioral outputs and overall organismal fitness [1-3]. At the neuronal level, the homeostatic response to stress depends on both synaptic and cell-intrinsic physiological processes that enable neurons to stably maintain optimal activity patterns [4-6]. Previous theoretical and empirical studies have suggested that the neuronal intrinsic robustness depends on the expression and activity of specific combinations of ion channels and transporters, which can vary across neuronal cell types and individuals [7-10]. While some of the transcriptional and physiological processes that enable neurons to adjust their intrinsic activity levels in response to long-term stressors have been identified [11-13], most of the genetic and molecular mechanisms that mediate susceptibility to acute, environmentally-induced seizures, such as fever-induced febrile seizures, remain unknown [14-16].

Because of its small size, large surface-to-volume ratio, and its inability to internally regulate body temperature, the fruit fly *Drosophila melanogaster*, represents an excellent model for studying mechanisms underlying the neuronal acute response to heat stress, which typically leads to seizure-like behavior, followed by paralysis [17-20]. Here we utilized this model to test the hypothesis that the knockdown of genes that are specifically important for the intrinsic neurophysiological homeostatic response to acute heat stress, would have little impact on fly behavior at permissive temperatures, but would lead to rapid paralysis under acute heat-stress conditions.

To test our hypothesis, we first employed a reverse genetic approach to identify candidate genes specifically involved in the neuronal homeostatic response to acute heat stress. By using a tissue-specific RNAi knockdown screen of voltage-gated potassium channels, we identified *seizure (sei)*, the fly ortholog of the mammalian hERG channel (KCNH2) [18, 21-25], as an essential element in the neuronal homeostatic response to acute heat stress. Previous studies have indicated that dominant hERG mutations, which are one of the primary genetic causes for the cardiac Long QT Syndrome (LQTS) in humans [26, 27], are also associated with a high prevalence of generalized seizures [28]. Yet, it is currently assumed that seizures in LQTS patients represent a derived secondary outcome of the primary cardiac pathology [29-31]. However, the data presented here, as well as previous studies that showed that ERG channels are expressed in mammalian neuronal tissues [32, 33], and contribute to intrinsic spike frequency adaptation in cultured mouse neuroblastoma cells and cerebellar Purkinje neurons [34, 35], suggest that ERG channels also have a specific function within the nervous system. By utilizing existing and novel genetic tools, here we show that the ERG channel *sei* is indeed essential for maintaining neuronal robustness under acute heat stress conditions in *Drosophila*. Specifically, we demonstrate that although *sei* is broadly expressed in the nervous system of the fly, its contribution to the organismal behavioral resistance to acute heat stress is primarily mediated via its specific action in excitatory cholinergic and glutamatergic neurons, the octopaminergic system, as well as non-neuronal glia. Furthermore, by generating a CRISPR/cas9-derived GFP-tagged allele of the native *sei* locus, we also show that at the subcellular level, *sei* exerts its action primarily in neuronal axons. Together, these studies indicate that mutations in hERG-like potassium channels may contribute directly to the etiology of stress-induced seizures in susceptible individuals by limiting the intrinsic neuronal homeostatic response to acute environmental stressors, possibly via homeostatic axonal spike frequency adaptation.

## Results

### Neuronal *sei* gene knockdown leads to acute heat-induced seizures

Previously published theoretical models and empirical studies have indicated that the actions of diverse voltage-gated potassium channels mediate action potential repolarization, and modulate the action potential threshold [36-39], which are important for the homeostatic regulation of synaptic activity and excitability [40-42]. Yet, which genes regulate the intrinsic capacity of neurons to buffer environmentally-induced hyperexcitability is mostly unknown. Thus, we initially hypothesized that the intrinsic ability of neurons to buffer acute heat stress is mediated, at least in part, by the action of specific voltage-gated potassium channels. Because most of these channels are expressed in both neuronal and non-neuronal tissues, we tested our hypothesis by using a neuronal-specific RNAi-dependent knockdown screen of all genes that encode voltage-gated potassium channels in the *Drosophila* genome. This screen revealed that neuronal knockdown of the gene *seizure (sei)*, which encodes the sole fly ortholog of hERG-like voltage-gated potassium channels [23, 24], has the strongest effect on lowering the threshold to heat-induced seizures (Fig 1A-B). Therefore, we focused our following primary working hypotheses on the contribution of the *sei* channel to the intrinsic neuronal homeostatic response to stress.

**Fig 1.**
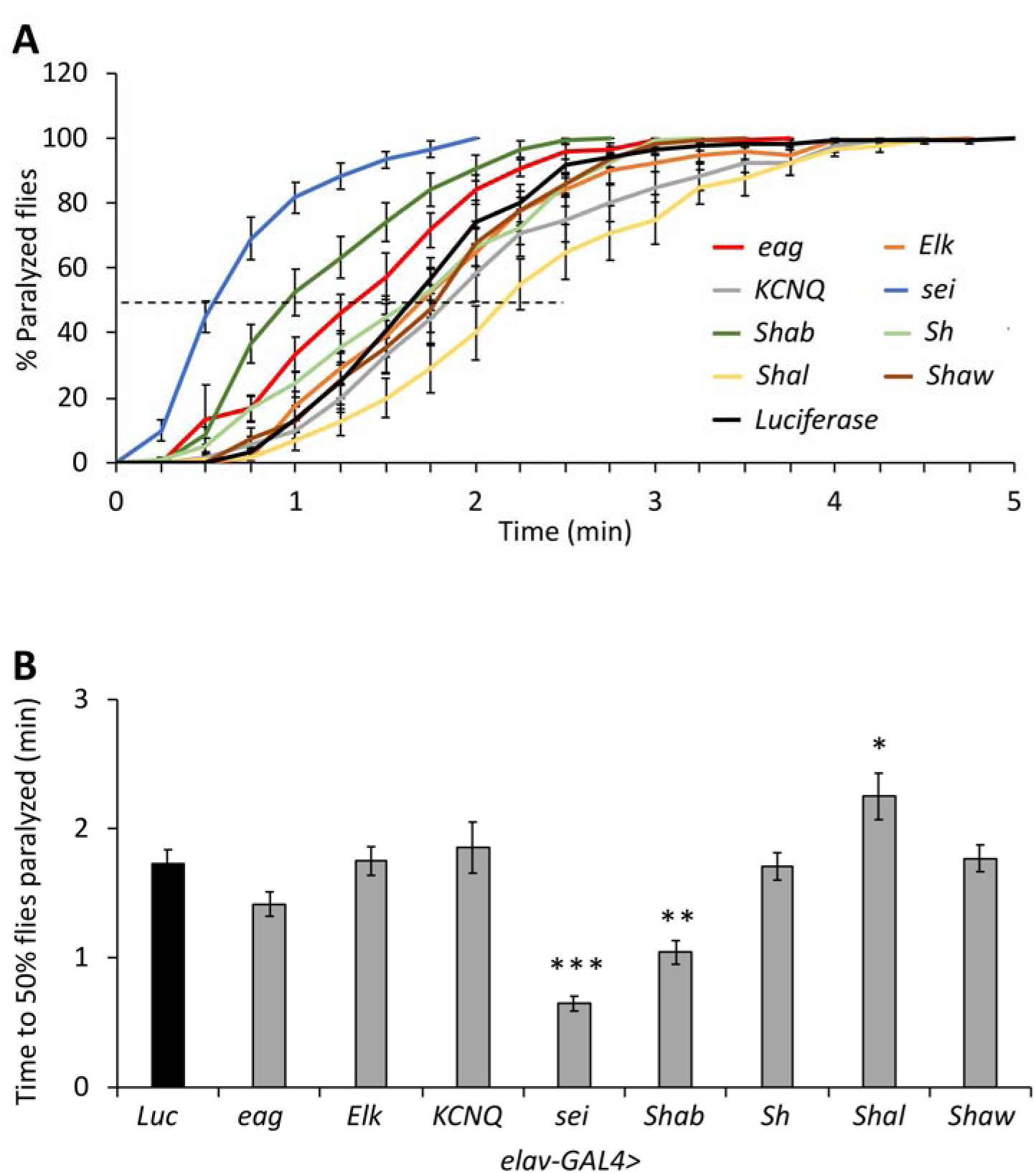
Neuronal (*elav-GAL4*) RNAi knockdown screen of *Drosophila* genes encoding voltage gated potassium channels. **A**) Cumulative percent paralyzed flies over time, with dotted line indicating 50% level. B) Direct comparison of time at which 50% of flies are paralyzed, using the same data as in A. n=12 vials/genotype. ANOVA (*p* < 0.0001) followed by Dunnett’s *post hoc* test was used to determine groups significantly different than the *Luciferase* control. Data are presented as mean ±SEM, \**p* < 0.05, \*\**p* < 0.01, \*\*\**p* < 0.001.

Mutations in the *seizure (sei)* gene were initially identified in a forward genetic screen for temperature-sensitive (ts) alleles of excitability-related genes in *Drosophila* [23, 24]. Because *sei* was assumed to be an essential component of general neuronal excitability, the two original EMS-induced *sei* mutant alleles were assumed to be structural ts alleles [18, 23-25]. However, because recent studies indicate that these *sei* alleles are more likely null or hypomorphic [25], we hypothesized that the action of the *sei* channel is specifically required for the ability of neurons to respond to acute heat stress. Thus, we next used a null allele of *sei* [43, 44] to demonstrate that *sei* activity is specifically required for the ability of adult flies to resist the impact of acute heat stress (Fig 2A-B). In addition to its role in the neuronal response to acute heat stress, we also show that the *sei* mutation impaired the ability of flies to adapt to a gradual heat stress (Fig 2C-D). Together, these data indicate that *sei* activity is not required for basal neuronal excitability, but is essential for the ability of neurons to maintain stable and adaptive firing rates under fluctuating environmental conditions.

**Fig 2.**
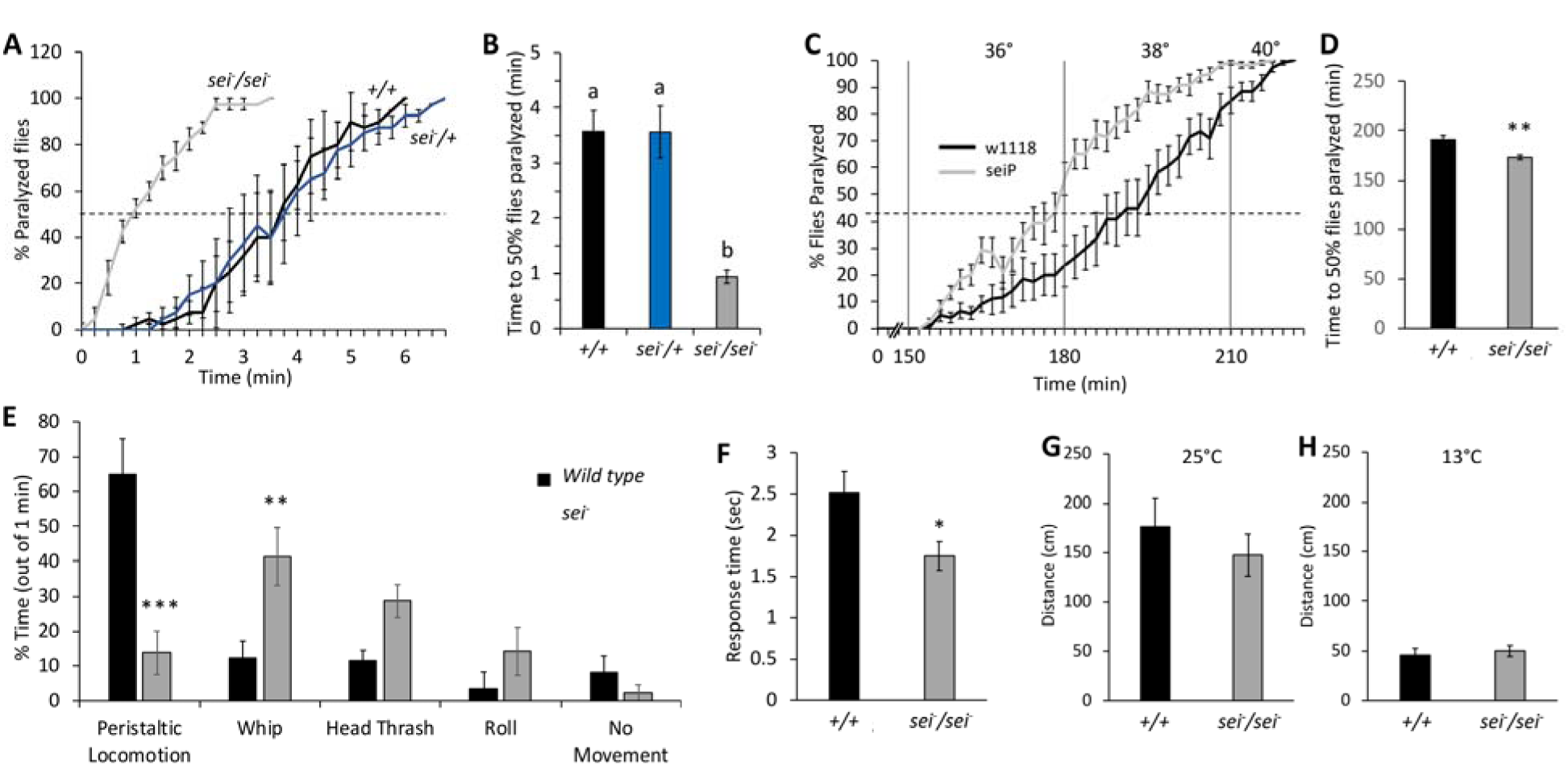
*sei* mutants display heat hypersensitivity. **A-B**) Acute heat assay paralysis behavior of homozygous and heterozygous *seiP* mutants and controls. n=4 vials/genotype. Letters above bars represent significantly different groups by ANOVA (*p* < 0.001) following Tukey’s *post hoc* analysis (*p* < 0.01). C-D) Gradual heat assay paralysis behavior of *seiP* mutant flies and controls. n=12 vials. E) Larval behavioral responses to heat in *seiP* mutants and controls. n=9. F) Response time of *seiP* mutant larvae and controls to thermal nociception assay. G-H) *seiP* mutant and wildtype larvae distance travelled in five minute trials at room temperature (25°; n=6) or cold temperature (13°; n=12). Data are presented as mean ±SEM, \**p* < 0.05, \*\**p* < 0.01, \*\*\**p* < 0.001 (Student’s *t*-test unless otherwise stated).

In contrast to adult flies, which under natural conditions could easily escape stressful environmental conditions by flying, larvae are much more constrained. Thus, the protective role of *sei* might be more ecologically relevant to the pre-adult developmental stages. Our data indicate that, as in the adult, *sei* activity is also necessary for normal larval locomotion under acute heat stress conditions (Fig 2E). Furthermore, because larvae can sense acute nociceptive stimuli, such as heat, via the activity of their cuticular multidendritic (md) sensory neurons [45-47], we next tested the hypothesis that heat induced hyperexcitability of md neurons in *sei* mutants would lead to nociception hypersensitivity. Indeed, we found that *sei* mutant larvae exhibit a significantly faster response to heat stimuli relative to wild type control (Fig 2F), which suggests that their nociceptive system is hypersensitive. Together, these studies indicate that the sei channel plays an important role in maintaining neuronal stability and robustness, and protecting *Drosophila* neurons from environmentally-induced hyperexcitability.

Because previous investigations of the impact of temperature changes on neuronal activity have shown that neurons will respectively increase or decrease their firing rates in response to a rise or fall in ambient temperature [48-50], we next hypothesized that *sei* mutant flies might be protected from the effect of acute cold stress on neuronal activity. However, we found no effect of the *s*ei mutation on larval locomotion at 13°C relative to wild type controls (Fig 2H). Thus, although the precise biophysical role of hERG-type voltage-gated potassium channels in regulating neuronal excitability remains elusive, the *in vivo* data presented here, as well as previously published *in vitro* studies [34, 35], indicate that hERG channels play a specific role in maintaining optimal neuronal activity by protecting neurons from environmentally-induced hyperexcitability but not hypoexcitability [25].

### Organismal resilience to heat stress requires the action of *sei* in excitatory and octopaminergic neurons, as well as glia

The *sei* gene is expressed in diverse neuronal and non-neuronal cell types, including cardiac and muscle cells [32, 51]. Previous studies by us and others have shown that mutations in *sei* increase the overall organismal sensitivity to acute heat stress [23-25]. Yet, whether this organismal phenotype is driven by the action of *sei* in all cell types that express it was unknown. Therefore, we next used tissue-specific RNAi knockdown to determine which cell types require *sei* activity to protect animals from heat-induced seizures. Similarly to our previous work [25], we found that neuronal-specific knockdown of *sei* is sufficient to phenocopy the effect of the null allele on the susceptibility of adult flies to heat-induced seizures (Fig 1A-B, 3A-B). However, we were also surprised to find that, although not as striking as in the neuronal knockdown, the organismal response to heat stress also depends on the activity of *sei* in non-neuronal glia (Fig 3C-D), but not in the heart or muscles (Fig 3E-H). Together, these data suggest that the organismal resistance to acute heat-induced seizures requires *sei* activity in the two primary cell types of the nervous system, neurons and glia.

**Fig 3.**
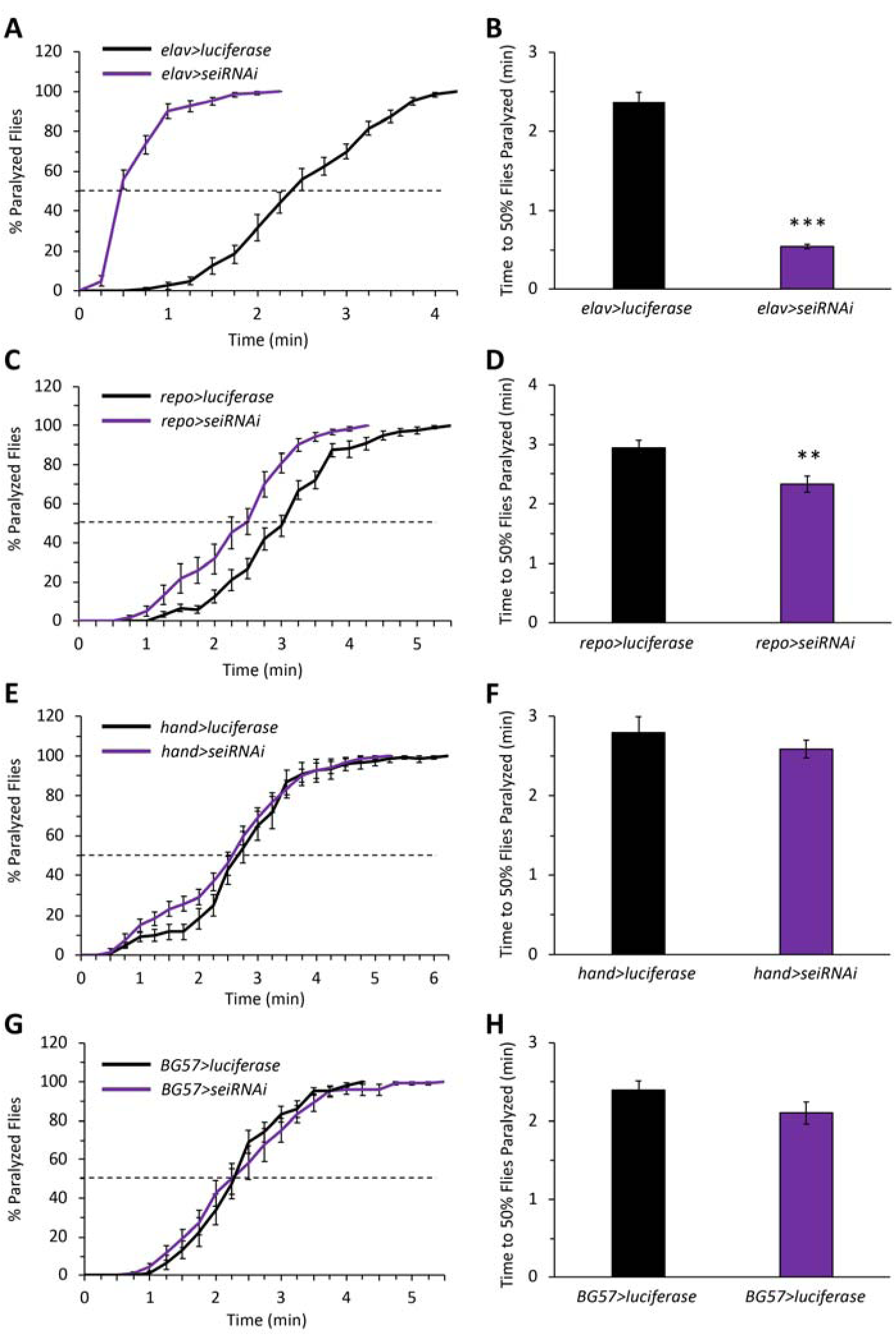
*sei* expression is required in neurons and glia for the homeostatic response to acute heat stress. A cell-type specific RNAi-dependent *sei* knockdown screen. A-B) Neurons (*elav-GAL4*); C-D) glia (*repo-GAL4*); E-F) heart (*hand-GAL4*); G-H) muscle (*BG57-GAL4*). n=12 vials/genotype. Data are presented as mean ±SEM, \*\**p* < 0.01, \*\*\**p* < 0.001 (Student’s *t*-test).

Because the fly CNS is a compact mosaic of different cell types, it is hard to establish whether *sei* is primarily expressed in neurons or glia. To address this, we generated a transgenic *Drosophila* driver line that expresses the *LexA* activator under the control of the putative *sei* promoter sequences [52, 53]. We then used this line to express a nuclear EGFP reporter, which indicated that *sei* is broadly expressed throughout the central nervous system (Fig 4A-B). To specifically identify *sei*-expressing glia, we next combined the *sei*-LexA line driving a nuclear EGFP with the glia-specific *Repo-GAL4* line driving a nuclear *RFP (RedStinger)*. Confocal imaging of co-labeled brains showed that in addition to its broad neuronal expression pattern, *sei* is also expressed in a small fraction of brain glia (Fig 4C-E, arrows). These data further indicate that the action of *sei* in the fly CNS is primarily mediated by its action in most neurons and few glia.

**Fig 4.**
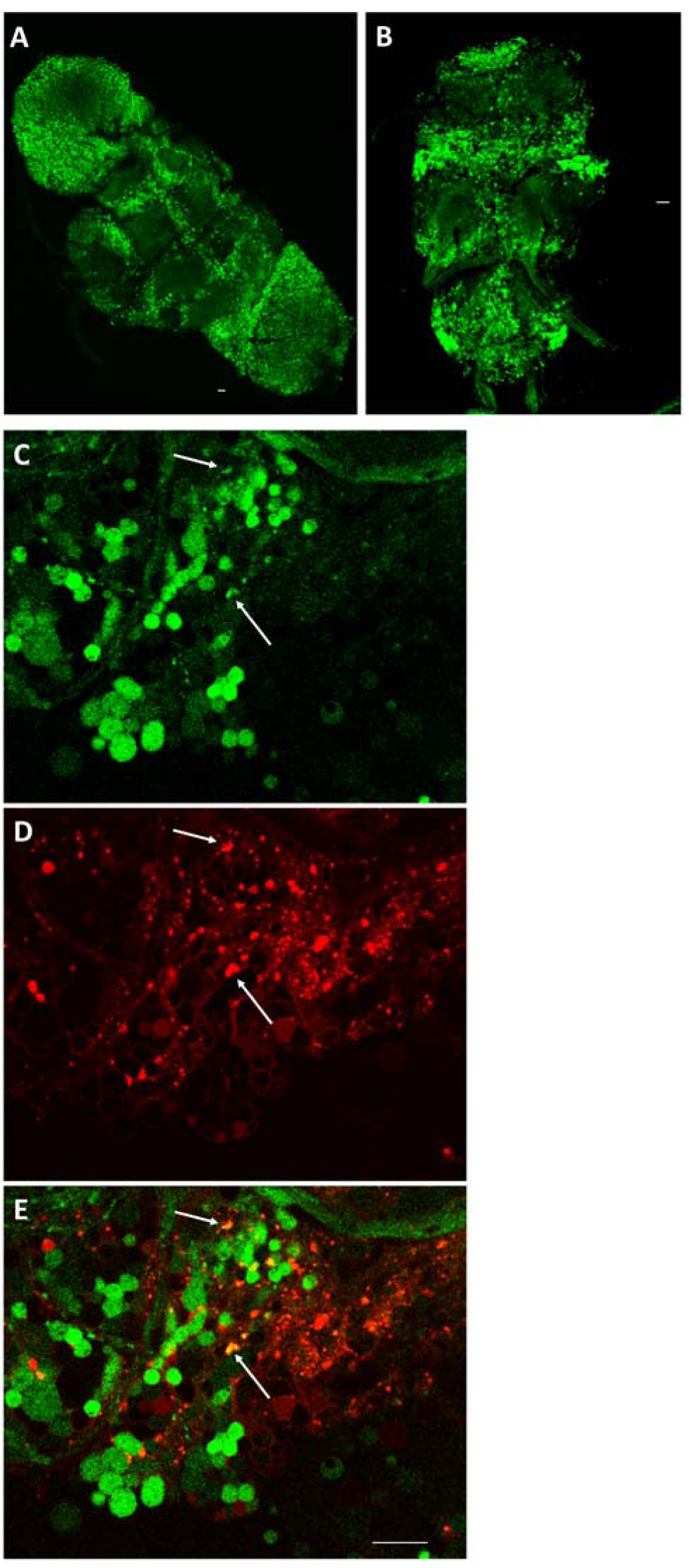
*sei* promoter drives expression in neurons and glia. **A-B**) Representative confocal z-stack images of an adult brain (A) and ventral nerve cord (B) from *sei-LexA>LexAOp-GFPnls* flies. C-E) Overlap of *sei-LexA>LexAOp-GFPnls* and glial marker *RepoGAL4>UAS-RedStinger*. Arrows point to examples of co-labeled cells.

Neurons are comprised of diverse cell types with different physiological properties and varying contributions to systems-level neural excitability. Therefore, we next wished to determine which neuronal subtypes might require sei activity for enabling the organismal response to acute heat stress. A broad screen of sei knockdown using several neuronal type-specific *GAL4* driver lines revealed that the organismal response to heat stress depends on the expression of *sei* in cholinergic (Fig 5A-B) and glutamatergic (Fig 5C-D) excitatory neurons, but not in GABAergic inhibitory neurons (Fig 5E-F). We also observed a significant effect of *sei* knockdown in the modulatory octopaminergic system (Fig 5G-H) but not in the dopaminergic, serotonergic, peptidergic, or the peripheral sensory systems (Fig 5I-P). Thus, our data suggest that *sei* plays an important role in protecting the nervous system from environmental stressors that could lead to general hyperexcitability and seizures by maintaining the neuronal robustness of excitatory and some neuromodulatory neurons.

**Fig 5.**
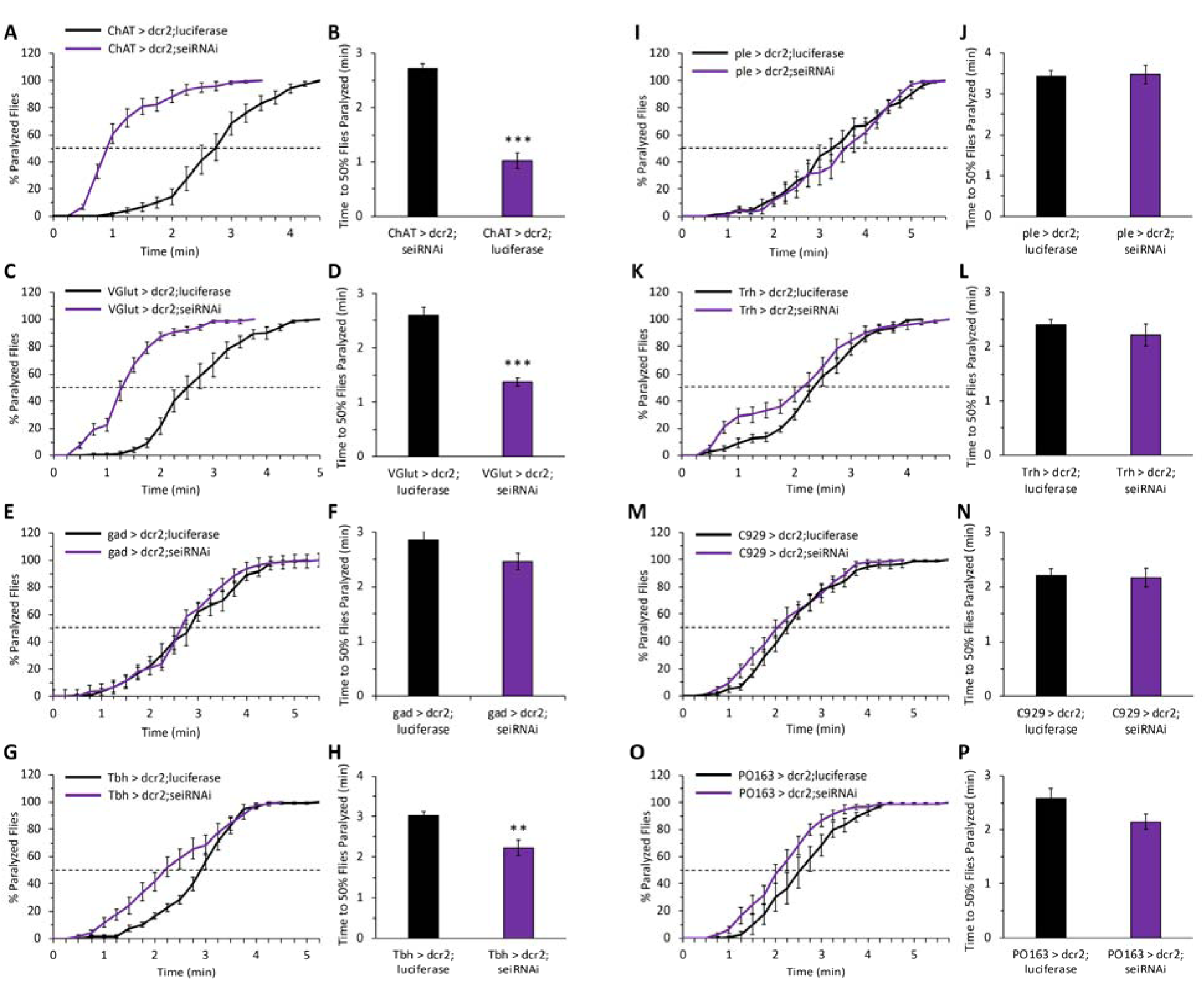
*sei* expression is specifically required in cholinergic, glutamatergic and octopaminergic cells for response to acute heat stress. A neuronal subtype specific RNAi-dependent sei knockdown screen. A-B) Cholinergic neurons (*ChAT-GAL4*) n=12 vials/genotype; C-D) Glutamatergic neurons (*gad-GAL4*) n=6; G-H) Octopaminergic neurons (*Tbh-GAL4*) n=12; E-F) GABAergic neurons (*Gad1-GAL4*) n=12; I-J) Dopaminergic neurons (*ple-GAL4*) n=12; K-L) Serotonergic neurons (*Trh-GAL4*) n=12; M-N) Peptidergic neurons (*C929-GAL4*) n=6; O-P) Sensory neurons (*PO163-GAL4*) n=6. Data are presented as mean ±SEM, \*\**p* < 0.01, \*\*\**p* < 0.001 (Student’s *t*-test).

### SEI channels are localized to axonal processes

The subcellular localization of various voltage gated ion channels plays an important role in determining how they might be contributing to neuronal signaling and excitability [54-56]. For example, ion channels localized to axons generally impact action potential generation, propagation, and modulation, while those localized to dendrites influence integration of synaptic inputs, propagation of electrical activity to the soma, and action potential backpropagation [54, 57, 58]. Nevertheless, ion channels with specific subcellular enrichment in either dendrites, cell bodies, axons, or presynaptic terminals have all been implicated in human epilepsies [59]. Therefore, we next determined the subcellular localization of native SEI channels by generating a C-terminus GFP-tagged allele of the endogenous *sei* locus (Fig 6A). We found that the response of homozygous *seiGFP* flies to heat stress is not different from wild type animals, which indicates that the tagged protein forms wild-type like channels (Fig 6B-C). We next used an anti-GFP antibody to probe the subcellular spatial distribution of *seiGFP* channels in the larval and adult nervous systems. These studies revealed that SEI is primarily localized to the axonal membranes of most neurons, but not in somas or dendrites (Fig 7). The enrichment of SEI channels in axons supports a model whereby ERG channels contribute to the intrinsic homeostatic regulation of optimal neuronal activity via the modulation of action potentials, a model further supported by the previously reported influence of mammalian ERG channels on spike frequency adaptation in cultured mammalian neurons [34, 35].

**Fig 6.**
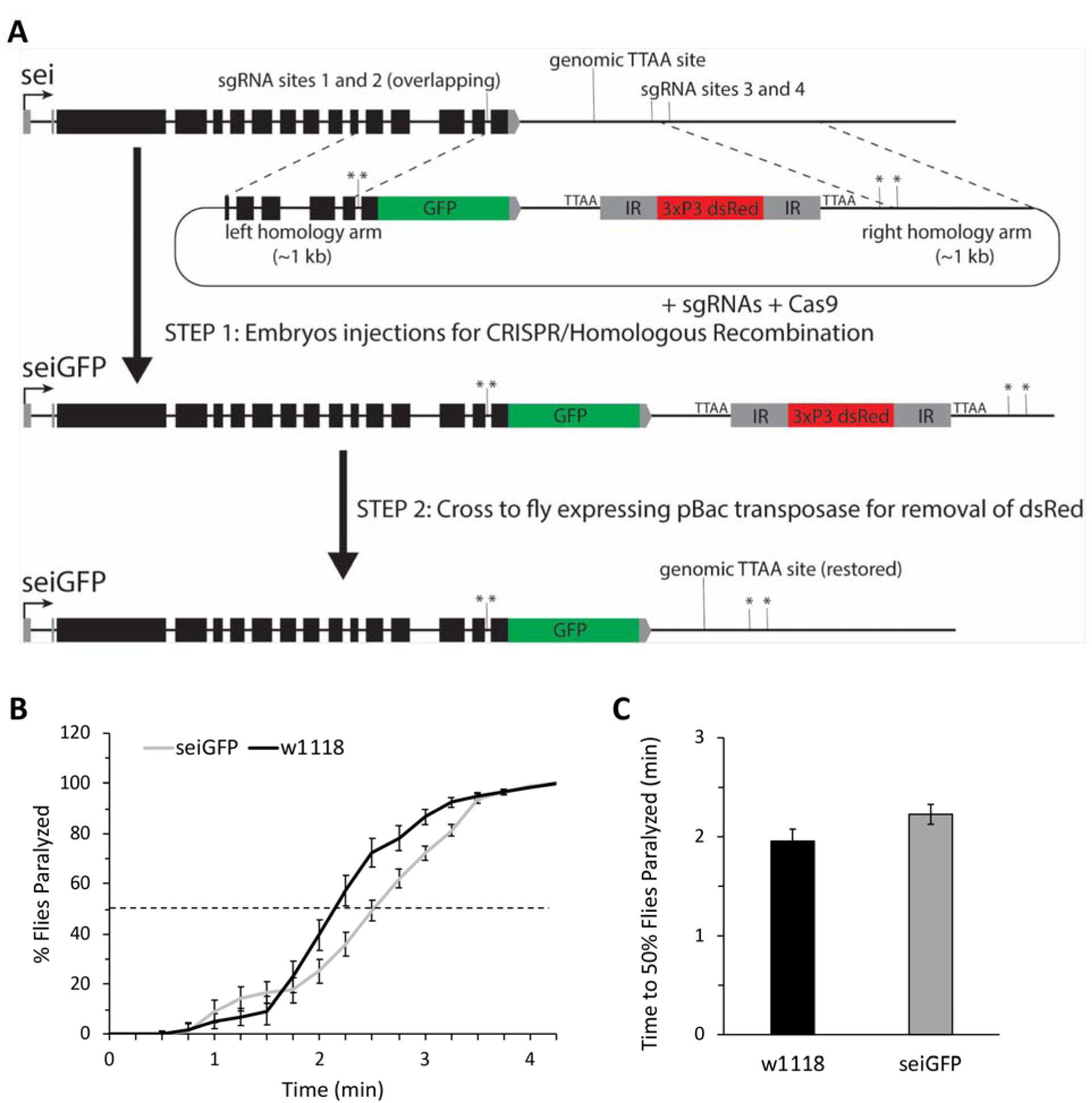
Generation and validation of the *seiGFP* allele. **A**) Strategy for the generation of the *seiGFP* allele by using CRISPR/*Cas9*-dependent DNA editing. Stars represent single nucleotide substitutions in the PAMs of sgRNA sites. B-C) Behavior of *seiGFP* and wildtype flies in the acute heat assay. n=12. Data was analyzed using Student’s *t*-test and presented as mean ±SEM.

**Fig 7.**
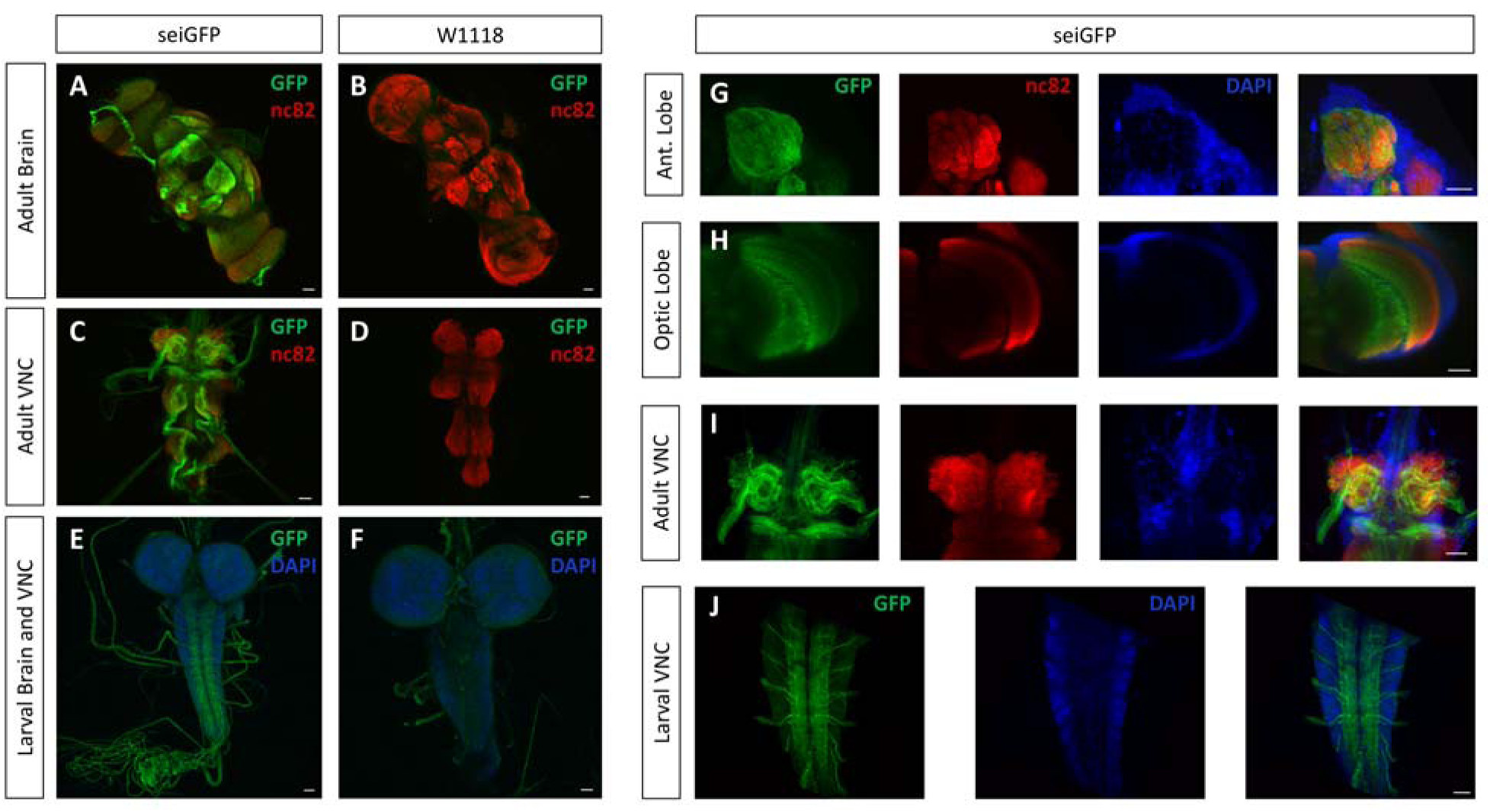
Nervous system expression of *seiGFP* in adult and larva is primarily localized to neuronal processes. **A-D**) Representative 20x confocal z-stack images of the adult brain (A-B) and adult ventral nerve cord (C-D) following immunostaining for *seiGFP* (green) and nc82 (red). E-F) 20x confocal z-stack images of the larval brain and ventral nerve cord of *seiGFP* (green) and DAPI (blue). G-I) 40x confocal z-stack images of the adult antennal lobe (G), optic lobe (H) and ventral nerve cord (I) with *seiGFP* (green), nc82 (red) and DAPI (blue). J) 40x confocal z-stack images of the larval ventral nerve cord of *seiGFP* (green) and DAPI (blue). Scalebars are 20µm.

## Discussion

Previous theoretical and empirical studies of neural circuit adaptability, and by extension, the ability of animals to maintain robust and adaptive behavioral outputs in unstable environments, depends on both the intrinsic homeostatic capacity of neurons to maintain an optimal activity pattern, and the ability of neural circuits to maintain stable outputs via the homeostatic regulation of neuronal connectivity and synaptic activity [4-6]. Yet, despite its high incidence, the majority of genetic and molecular factors that regulate neuronal homeostasis, and increase susceptibility to seizures, remain mostly unknown [14-16]. Here, we show that the *Drosophila* voltage-gated potassium channel *sei*, a conserved ortholog of the human hERG gene, is required for the neuronal homeostatic response to acute heat stress. Furthermore, we show that the organismal capacity to buffer the effects of acute heat stress depends on the independent activity of *sei* in both neurons and glia. We also found that although *sei* is broadly expressed in the nervous system, its contribution to the overall organismal resistance to acute heat stress seems to be specifically driven by its action in cholinergic and glutamatergic excitatory neurons, as well as to a lesser extent in the modulatory octopaminergic system. In addition, we developed genetic tools to show that SEI is expressed in the axons of neurons and in glia. Together, our data highlight the important role of *sei* in the organismal homeostatic response to acute environmental stress, by providing robustness to both the intrinsic activity of specific neuronal populations, and the neural circuits that harbor them.

How ERG-type potassium channels might contribute to neuronal intrinsic homeostasis during bouts of acute stress is not well understood. Nonetheless, *in vivo* and *in vitro* studies in *Drosophila* and mammalian models have suggested that ERG channels have little effect on baseline neuronal firing rate, but can prevent rapid firing in response to environmental or electrophysiological stimuli that induce hyperexcitability [25, 34, 35]. We have previously shown that in *Drosophila* motor neurons, basal neuronal firing patterns are unaffected by the sei mutation at optimal 25°C, but become hyperexcitable in response to a rapid temperature increase [25]. Similarly, in electrophysiological studies of mammalian brain slices, *in vitro* cultured neurons, and heterologously-expressed mammalian ERG channels, pharmacological blockers of hERG channels have little effect on firing rates in response to small current injections, but greatly diminish spike frequency adaptation in response to large current injections, resulting in rapid firing rates [34, 35]. The presence of SEI channels specifically in axons (Fig 7) suggests that they do not affect the propagation of dendritic potentials, but rather limit the rate of action potential generation and propagation, which is sufficient to prevent rapid firing rates. Together, these data suggest a model whereby ERG-like potassium channels play a crucial role in mediating the neuronal homeostatic response to acute stress by protecting neurons from rapid increase in firing rates, and therefore, support neuronal robustness when exposed to extreme environmental fluctuations.

At the neuronal network level, seizures are thought to result from an imbalance between excitatory and inhibitory neural signaling pathways [15, 60]. We found that knocking down *sei* specifically in all cholinergic neurons, the primary excitatory pathway in the fly central nervous system, is sufficient to phenocopy the effects of *sei* null mutations on the organismal resistance to heat-induced seizures. These results are similar to previous studies, which showed that increasing activity of the cholinergic system in flies, via genetic manipulations of voltage gated sodium channels and optogenetic neural activation, is sufficient to increase seizure-related and paralytic behavior [61-64]. The simplest interpretation of these data together is that the lack of *sei* in cholinergic excitatory neurons makes them hypersensitive to heat-induced hyperexcitability, which subsequently surpasses the buffering capacity of the inhibitory neurotransmission pathways, and therefore leads to the rapid development of generalized seizures and paralysis.

Additionally, we observed a large increase in seizure susceptibility when *sei* is knocked down in glutamatergic neurons. Although we and others have previously demonstrated that the activity of motor neurons, which in insects are primarily glutamatergic, is increased in seizure-susceptible mutant flies [61, 65], this finding suggests that the decreased intrinsic ability of motor neurons to resist acute stress is sufficient for inducing organismal seizure-like phenotype (Fig 5C-D). Nevertheless, we currently cannot exclude the possibility that the observed effect of knocking down *sei* expression in glutamatergic neurons on organismal sensitivity to stress is mediated via the action of a small number of modulatory glutamatergic neurons within the central nervous system [66].

We also observed an impairment in the organismal homeostatic response to acute heat stress when *sei* is specifically knocked-down in the modulatory octopaminergic system (Fig 5G-H). Previous studies of the octopaminergic system in *Drosophila* and other insects have indicated that octopamine and related biogenic amines have broad impact on diverse neuronal processes at the developmental and physiological timescales [67, 68]. Because *sei* mutant flies seem to have normal behaviors when housed under constant optimal conditions, it is likely that the effects of knocking down sei in octopaminergic neurons on heat-induced seizures are physiological, not developmental. Although we currently do not know which specific elements of the octopaminergic system play a role in the organismal response to acute heat stress, previous work has shown that exogenous application of octopamine in *Drosophila* increases contraction force of muscles and their response to synaptically driven contractions [69]. Therefore, one possible mechanism by which *sei* knockdown in octopaminergic neurons might affect observed heat-induced seizures is via the direct modulation of the neuromuscular junction. Octopamine has also been shown to play important roles in the central nervous system, including modulation of behaviors related to motivation, sleep, aggression, social behaviors and learning and memory [67, 70-73]. Therefore, *sei* knockdown in octopaminergic neurons may result in a broader shift in synaptic processes associated with the homeostatic maintenance of the balance between excitatory and inhibitory pathways under acute stress conditions.

The important role of *sei* activity in regulating the capacity of the nervous system to buffer acute environmental stress is further supported by our discovery that its knockdown in glia also increased susceptibility to acute heat-induced seizures (Fig 3C-D). Previous studies have suggested that some glia cells are important for maintaining synaptic activity and homeostasis, and that disrupting glia functions could contribute to the etiology of seizures because of their role in modulating extracellular potassium concentration, adenosine levels, the size of the extracellular space, and uptake of neurotransmitters [74-79]. Additionally, glia in the mammalian hippocampus express muscarinic and nicotinic acetylcholine receptors [80-82], and acetylcholine release in the hippocampus has been shown to increase GABAergic signaling through activation of glial cells, possibly by glial release of glutamate [83]. If conserved, this would be an intriguing mechanism whereby glia might control excitability throughout, or in specific regions of, the *Drosophila* central nervous system. However, how the action of voltage gated ion channels such as *sei* in glia might affect these specific processes remains mostly unknown. Nevertheless, glial expression of another voltage gated potassium channel that is associated with human epilepsy, KCNJ10, has been shown to lead to epileptic activity in a mouse model, possibly via its role in buffering extracellular potassium and glutamate [84, 85]. Whether hERG-like channels play a similar role in glia remains to be explored.

Together, the data we present here provide important insights into the possible role of hERG channels in regulating neuronal robustness and susceptibility to stress-induced seizures. From a clinical perspective, our data suggest that the high incidence of generalized seizures that has been reported in LQTS patients that carry mutations in the hERG genes [28] might not be a secondary cardiogenic comorbidity, as is currently often assumed [29-31]. Instead, because about 40% of patients that carry LQTS-related hERG mutations have reported a personal history of seizures, as compared to less than 20% in LQTS patients with similar cardiac pathologies that are due to mutations in other genes [28], we hypothesize that seizure etiology in many LQTS patients is likely due to the direct impact of mutations in hERG on nervous system functions, independent of their cardiovascular condition. Therefore, we predict that it is possible that some unidentified mutations in hERG might be causally related to epilepsies, independent of the presentation of any LQTS-related pathologies, and may represent novel genetic risk factors for seizures.

The studies we describe here provide compelling evidence that hERG channels play an essential role in protecting the nervous system from acute environmental stressors, such as heat, which could potentially lead to hyperexcitability and seizures. Furthermore, we show that in *Drosophila*, the activity of the ERG channel *sei* contributes to neuronal and behavioral robustness via its action in independent cell types in the nervous system. These important insights should help us to better understand how the nervous system responds to acute environmental stressors, and possibly provide important mechanistic insights into some of the known pathologies associated with hERG mutations in human patients.

## Materials and Methods

### Fly Stocks and Genetics

Flies (*Drosophila melanogaster*) were raised on standard corn syrup-soy food (Archon Scientific) at 25°C temperature, 70% humidity, on a 12:12 light/dark cycle. Unless specifically noted, wild type control line used was *w*^*1118*^. All fly strains were either produced in the Ben-Shahar lab or obtained from the Bloomington Stock Center (stock numbers in parentheses). UAS-RNAi TRiP lines [86] used in the initial screen included *sei* (#31681), *shab* (#25805), *eag* (#31678), *shaker* (#53347), *shaw* (#28346), *shal* (#31879), *elk* (#25821) and *kcnq* (#27252). The TRiP UAS-*Luciferase* RNAi was used a control (#35789), and UAS-RNAi lines were driven by the *elav-GAL4*; *UAS-Dicer2* line (#25750) (Fig 1A-B). For the cell-type-specific *sei* knockdown screen (Figs 3 and 5), the UAS-RNAi for sei and *luciferase* were each recombined with *UAS-Dcr2*. The original null *seiP* allele from Bloomington (#21935) was backcrossed into the *w*^*1118*^ wild type strain. Other transgenic lines from the Bloomington stock center included: *UAS-RedStinger* (#8546), *LexAOp-GFPnls* (#29954), *elav-GAL4* [87] (#458), *Repo-GAL4* (#7415), *hand-GAL4* (#48396), *ChAT-GAL4* (#6798), *VGlut-GAL4* (#60312), *gad-GAL4* (#51630), *ple-GAL4* (#8848), *Tbh-GAL4* (#39939), *Trh-GAL4* (#49258), and *C929-GAL4* (#25373). *BG57-GAL4* [88] and *PO163-GAL4* were from the Dickman and Zlatic labs respectively.

The C-terminus GFP-tagged allele of *sei* was generated via CRISPR/*Cas9*-dependent editing by using a modified “scarless” strategy (www.flyCRISPR.molbio.wisc.edu)[89, 90]. Specifically, four sgRNAs TGTAAGCGAATACCACGTTG, GACAGCATTCTCCCGCAACG, GAAGCAGAAGCAGGTAACTC, AGGTGAGTGAGTTACTCATC, which flank the targeted genomic sei sequence were designed using flyRNAi.org/crispr. Complementary oligos that correspond to each individual sgRNA (IDT) were cloned into the pDCC6 plasmid (a gift from Peter Duchek, Addgene plasmid # 59985), which also includes the coding sequence for *Cas9* [91], by using the BbsI restriction enzyme (NEB). The donor plasmid for homologous recombination was constructed by using a Golden Gate assembly [92] to recombine four DNA elements: 1) A backbone with ampicillin resistance (pBS-GGAC-ATGC plasmid (a gift from Frank Schnorrer, Addgene #60949)[93]; 2) Eye-specific dsRed reporter driven by the 3XP3 promoter, flanked by two *PiggyBac* transposase recognition sites, which was PCR-amplified from the pHD-ScalessDsRed plasmid (a gift from Kate O’Connor-Giles, Drosophila Genome Resource Center #1364) using the following primers 5’-CACACCACGTCTCATTAACCCTAGAAAGATAATCATATTGTG-3’ and 5’-CACACCACGTCTCACCCTAGAAAGATAGTCTGCGT-3’ (the primers included BsmBI restriction enzyme sites and overhangs corresponding to the left and right homology arms); 3-4) The left and right homology arms, which consisted of 1kb genomic DNA fragments upstream and downstream of the sgRNA sites respectively. Single base pair mutations were introduced into the PAM sequence at each sgRNA binding site on the homology arms to prevent Cas9-dependent cutting of the donor plasmid. The right homology arm also included the GFP coding sequence in frame with the 3’ end of the last coding exon of sei, immediately upstream of the endogenous stop codon (Fig 6A). pDCC6 plasmids containing sgRNA and cas9 sequences (100 ng/µL) and the donor plasmid (500 ng/µL) were co-injected into *y*^-^*w*^-^ background. Subsequently, correct genomic integration of the GFP tag was verified by screening for DsRed-positive animals, followed by sequencing of genomic PCR fragments. The final tagged *sei* allele was generated by removing the DsRed cassette via the introduction of the *piggyBac* transposase (Bloomington #8285) (Fig 6A).

The *sei-LexA* transgenic flies were generated by amplifying a 2612 bp genomic DNA fragment upstream of the *sei* start codon by using the following PCR primers: 5’-GTCGACCGCCGGCAAAGTATCAACAT-3’ and 5’-GCGGCCGCTTTTAAGTCTGCAAAGTATAGAAACG-3’, followed by cloning into the pENTR1A plasmid (ThermoFisher) with SalI and NotI restriction enzymes. The *sei* promoter fragment was then recombined into the pBPnlsLexA::p65Uw vector (a gift from Gerald Rubin, Addgene plasmid # 26230)[52] by using the Gateway reaction (ThermoFisher). The *sei-LexA* containing plasmid was integrated into the fly genome by using a line carrying a PhiC31 integrase landing site on Chromosome III (Bloomington #24483)[94].

### Behavioral response to acute heat stress

Assays were performed as previously described [25]. In short, two-day old flies were collected and transferred into standard vials containing food (five per sex in each vial). On the following day, flies were flipped into an empty vial, and tested within the next hour. For testing, vials with 10 flies were individually submerged into a 41-42°C water bath and observed for seizure-like behavior and paralysis. The cumulative number of paralyzed flies (immobile at the bottom of the vial) was recorded every 15 seconds. The time at which 50% of the flies in a given vial were paralyzed was used as a measure of seizure susceptibility for statistical analysis.

### Behavioral response to gradual heat stress

Two-day old flies were housed in groups of 10 as above. Subsequently, vials were placed in a temperature-controlled incubator (Fisher Scientific Isotemp) with a glass door, which allowed continuous video recording of their behavior. To test for the ability of flies to adapt to gradual temperature increase, flies were first acclimated to 26°C followed by a 2°C increase every 30 minutes to a maximum of 42°C. The number of paralyzed flies was recorded every two minutes.

### Larval Locomotion Behavior

A 60mm plastic petri dish was filled with 3% agar, and placed on a Peltier plate surface of a PCR machine, which was set to either 37°C or 13°C for the heat or cold stress respectively. Identical tests at 25°C were used as controls. For the heat stress condition, locomotion was assayed by placing individual third instar foraging larvae on the agar surface and video recording (Logitech C920 Webcam) for one minute. Larva locomotion was analyzed by assessing the amount of time spent executing the following specific behaviors: peristaltic locomotion, whipping, head thrashing, rolling and no movement [95](S1 Video). For the cold stress condition, individual larvae were allowed to acclimate for 30s before their behavior was recorded for four min. Behaviors of all larvae were tracked and analyzed by using a custom designed motion tracker system (McKinney and Ben-Shahar, unpublished), which enabled computer-assisted analyses of distance traveled.

### Larval nociceptive response to heat

Foraging 3^rd^ instar larvae were removed from bottles, washed in water, and placed on a water-saturated 3% agar plate. Behavioral tests were conducted in constant 25°C and 70% humidity. Larvae were allowed to acclimate to the agar plate for 10 seconds. A custom-made heat probe (Thermal Solutions Controls and Indicators Corporation) set to 50°C was used to gently touch the side of the larva’s body, and the amount of time for the larva to roll over was recorded. For each genotype, 47-50 larvae were individually tested.

### Immunostaining and imaging

Third-instar larval brains were dissected and fixed in 4% PFA for 20 minutes, washed with PBST+0.1% Triton-X (PBST), and then blocked with Superblock (ThermoFisher) for one hour. Larval brains were subsequently incubated overnight at 4°C with rabbit anti-GFP (A-11122, ThermoFisher) diluted in Superblock at 1:1000. After washing with PBST, larval brains were incubated for 2 hours at room temperature with secondary antibody Alexa Fluor 488-conjugated goat anti-rabbit (A-11034, ThermoFisher) and FITC-conjugated goat anti-HRP (123-095-021, Jackson ImmunoResearch) to label neurons, each diluted 1:1000 in Superblock. After secondary incubation, brains were washed in PBST, mounted using FluoroGel II with DAPI (ThermoFisher), and imaged using an A1Si laser scanning confocal microscope (Nikon). Adult brains were dissected, fixed and blocked as above, then subsequently incubated overnight at 4°C in rabbit anti-GFP (A-11122, ThermoFisher) diluted at 1:1000 and mouse anti-Brp (NC82; Developmental Studies Hybridoma Bank) diluted 1:33 in Superblock. After washing with PBST, adult brains were incubated overnight at 4°C with secondary antibodies Alexa Fluor 488-conjugated anti-rabbit (A-11034, ThermoFisher) and Alexa Fluor rhodamine-conjugated donkey anti-mouse (sc-2300, Santa Cruz Biotechnology), each diluted 1:500 in Superblock. Adult brains were mounted and imaged as above. To image live brains expressing either *RedStinger* or *GFPnls* nuclear markers, the tissues were dissected, mounted in PBS, and imaged on a confocal microscope within one hour of the dissection.

### Statistical analysis

Indicated statistical comparisons were analyzed by using Excel (Microsoft) and Prism 7 (GraphPad). Statistical significance was set at *p*<0.05.

## Acknowledgements

We thank the Bloomington Drosophila Stock Center (NIH P40OD018537) and the TRiP at Harvard Medical School (NIH/NIGMS R01-GM084947) for providing transgenic fly stocks, and the Drosophila Genome Resource Center (NIH grant 2P40OD010949) for plasmids used in these studies.

## Supporting Information

**S1 Video. Heat induced *seiP* mutant larval behavior.**

## References

1. Kagias K, Nehammer C, Pocock R. Neuronal responses to physiological stress. Front Genet. 2012;3:222. Epub 2012/11/01. doi: 10.3389/fgene.2012.00222. PubMed PMID: 23112806; PubMed Central PMCID: PMCPMC3481051.

2. Davis GW. Homeostatic signaling and the stabilization of neural function. Neuron. 2013;80(3):718–28. Epub 2013/11/05. doi: 10.1016/j.neuron.2013.09.044. PubMed PMID: 24183022; PubMed Central PMCID: PMCPMC3856728.

3. Ramocki MB, Zoghbi HY. Failure of neuronal homeostasis results in common neuropsychiatric phenotypes. Nature. 2008;455(7215):912–8. doi: 10.1038/nature07457. PubMed PMID: 18923513; PubMed Central PMCID: PMC2696622.

4. Marder E, Goaillard JM. Variability, compensation and homeostasis in neuron and network function. Nat Rev Neurosci. 2006;7(7):563–74. doi: 10.1038/nrn1949. PubMed PMID: 16791145.

5. Debanne D, Poo MM. Spike-timing dependent plasticity beyond synapse - pre- and post-synaptic plasticity of intrinsic neuronal excitability. Front Synaptic Neurosci. 2010;2:21. doi: 10.3389/fnsyn.2010.00021. PubMed PMID: 21423507; PubMed Central PMCID: PMCPMC3059692.

6. Turrigiano G. Too many cooks? Intrinsic and synaptic homeostatic mechanisms in cortical circuit refinement. Annu Rev Neurosci. 2011;34:89–103. doi: 10.1146/annurev-neuro-060909-153238. PubMed PMID: 21438687.

7. O’Leary T, Williams AH, Caplan JS, Marder E. Correlations in ion channel expression emerge from homeostatic tuning rules. Proc Natl Acad Sci U S A. 2013;110(28):E2645–54. doi: 10.1073/pnas.1309966110. PubMed PMID: 23798391; PubMed Central PMCID: PMCPMC3710808.

8. Drion G, O’Leary T, Marder E. Ion channel degeneracy enables robust and tunable neuronal firing rates. Proc Natl Acad Sci U S A. 2015;112(38):E5361–70. doi: 10.1073/pnas.1516400112. PubMed PMID: 26354124; PubMed Central PMCID: PMCPMC4586887.

9. Schulz DJ, Goaillard JM, Marder E. Variable channel expression in identified single and electrically coupled neurons in different animals. Nat Neurosci. 2006;9(3):356–62. doi: 10.1038/nn1639. PubMed PMID: 16444270.

10. Marder E. Variability, compensation, and modulation in neurons and circuits. Proc Natl Acad Sci U S A. 2011;108 Suppl 3:15542–8. doi: 10.1073/pnas.1010674108. PubMed PMID: 21383190; PubMed Central PMCID: PMCPMC3176600.

11. Desai NS, Rutherford LC, Turrigiano GG. Plasticity in the intrinsic excitability of cortical pyramidal neurons. Nat Neurosci. 1999;2(6):515–20. doi: 10.1038/9165. PubMed PMID: 10448215.

12. Golowasch J, Abbott LF, Marder E. Activity-dependent regulation of potassium currents in an identified neuron of the stomatogastric ganglion of the crab Cancer borealis. J Neurosci. 1999;19(20):RC33. PubMed PMID: 10516335.

13. Ransdell JL, Nair SS, Schulz DJ. Rapid homeostatic plasticity of intrinsic excitability in a central pattern generator network stabilizes functional neural network output. J Neurosci. 2012;32(28):9649–58. doi: 10.1523/JNEUROSCI.1945-12.2012. PubMed PMID: 22787050.

14. Ottman R, Risch N. Genetic Epidemiology and Gene Discovery in Epilepsy. In: Noebels JL, Avoli M, Rogawski MA, Olsen RW, Delgado-Escueta AV, editors. Jasper’s Basic Mechanisms of the Epilepsies. 4th ed. Bethesda (MD)2012.

15. Staley K. Molecular mechanisms of epilepsy. Nat Neurosci. 2015;18(3):367–72. doi: 10.1038/nn.3947. PubMed PMID: 25710839; PubMed Central PMCID: PMCPMC4409128.

16. Feng B, Chen Z. Generation of Febrile Seizures and Subsequent Epileptogenesis. Neurosci Bull. 2016;32(5):481–92. Epub 2016/08/27. doi: 10.1007/s12264-016-0054-5. PubMed PMID: 27562688; PubMed Central PMCID: PMCPMC5563761.

17. Burg MG, Wu CF. Mechanical and temperature stressor-induced seizure-and-paralysis behaviors in Drosophila bang-sensitive mutants. J Neurogenet. 2012;26(2):189–97. Epub 2012/06/22. doi: 10.3109/01677063.2012.690011. PubMed PMID: 22716921; PubMed Central PMCID: PMCPMC3398232.

18. Kasbekar DP, Nelson JC, Hall LM. Enhancer of seizure: a new genetic locus in Drosophila melanogaster defined by interactions with temperature-sensitive paralytic mutations. Genetics. 1987;116(3):423–31. PubMed PMID: 2440763; PubMed Central PMCID: PMCPMC1203154.

19. Abram PK, Boivin G, Moiroux J, Brodeur J. Behavioural effects of temperature on ectothermic animals: unifying thermal physiology and behavioural plasticity. Biol Rev Camb Philos Soc. 2017;92(4):1859–76. Epub 2017/10/06. doi: 10.1111/brv.12312. PubMed PMID: 28980433.

20. Neven LG. Physiological responses of insects to heat. Postharvest Biol Tec. 2000;21(1):103–11. doi: Doi 10.1016/S0925-5214(00)00169-1. PubMed PMID: WOS:000166230900010.

21. Jackson FR, Wilson SD, Strichartz GR, Hall LM. Two types of mutants affecting voltage-sensitive sodium channels in Drosophila melanogaster. Nature. 1984;308(5955):189–91. PubMed PMID: 6322008.

22. Jackson FR, Gitschier J, Strichartz GR, Hall LM. Genetic modifications of voltage-sensitive sodium channels in Drosophila: gene dosage studies of the seizure locus. J Neurosci. 1985;5(5):1144–51. PubMed PMID: 2582101.

23. Titus SA, Warmke JW, Ganetzky B. The Drosophila erg K+ channel polypeptide is encoded by the seizure locus. J Neurosci. 1997;17(3):875–81. PubMed PMID: 8994042.

24. Wang XJ, Reynolds ER, Deak P, Hall LM. The seizure locus encodes the Drosophila homolog of the HERG potassium channel. J Neurosci. 1997;17(3):882–90. PubMed PMID: 8994043.

25. Zheng X, Valakh V, Diantonio A, Ben-Shahar Y. Natural antisense transcripts regulate the neuronal stress response and excitability. Elife. 2014;3:e01849. doi: 10.7554/eLife.01849. PubMed PMID: 24642409; PubMed Central PMCID: PMC3953951.

26. Curran ME, Splawski I, Timothy KW, Vincent GM, Green ED, Keating MT. A molecular basis for cardiac arrhythmia: HERG mutations cause long QT syndrome. Cell. 1995;80(5):795–803. PubMed PMID: 7889573.

27. Schwartz PJ, Crotti L, Insolia R. Long-QT syndrome: from genetics to management. Circ Arrhythm Electrophysiol. 2012;5(4):868–77. doi: 10.1161/CIRCEP.111.962019. PubMed PMID: 22895603; PubMed Central PMCID: PMCPMC3461497.

28. Johnson JN, Hofman N, Haglund CM, Cascino GD, Wilde AA, Ackerman MJ. Identification of a possible pathogenic link between congenital long QT syndrome and epilepsy. Neurology. 2009;72(3):224–31. doi: 10.1212/01.wnl.0000335760.02995.ca. PubMed PMID: 19038855; PubMed Central PMCID: PMCPMC2677528.

29. MacCormick JM, McAlister H, Crawford J, French JK, Crozier I, Shelling AN, et al. Misdiagnosis of long QT syndrome as epilepsy at first presentation. Ann Emerg Med. 2009;54(1):26–32. doi: 10.1016/j.annemergmed.2009.01.031. PubMed PMID: 19282063.

30. Zaidi A, Clough P, Cooper P, Scheepers B, Fitzpatrick AP. Misdiagnosis of epilepsy: many seizure-like attacks have a cardiovascular cause. J Am Coll Cardiol. 2000;36(1):181–4. PubMed PMID: 10898432.

31. Hunt DP, Tang K. Long QT syndrome presenting as epileptic seizures in an adult. Emerg Med J. 2005;22(8):600–1. doi: 10.1136/emj.2003.007997. PubMed PMID: 16046776; PubMed Central PMCID: PMC1726881.

32. Wymore RS, Gintant GA, Wymore RT, Dixon JE, McKinnon D, Cohen IS. Tissue and species distribution of mRNA for the IKr-like K+ channel, erg. Circ Res. 1997;80(2):261–8. PubMed PMID: 9012748.

33. Warmke JW, Ganetzky B. A family of potassium channel genes related to eag in Drosophila and mammals. Proc Natl Acad Sci U S A. 1994;91(8):3438–42. PubMed PMID: 8159766; PubMed Central PMCID: PMCPMC43592.

34. Chiesa N, Rosati B, Arcangeli A, Olivotto M, Wanke E. A novel role for HERG K+ channels: spike-frequency adaptation. J Physiol. 1997;501 (Pt 2):313–8. PubMed PMID: 9192303; PubMed Central PMCID: PMC1159479.

35. Sacco T, Bruno A, Wanke E, Tempia F. Functional roles of an ERG current isolated in cerebellar Purkinje neurons. J Neurophysiol. 2003;90(3):1817–28. doi: 10.1152/jn.00104.2003. PubMed PMID: 12750425.

36. Byrne JH. Analysis of ionic conductance mechanisms in motor cells mediating inking behavior in Aplysia californica. J Neurophysiol. 1980;43(3):630–50. Epub 1980/03/01. doi: 10.1152/jn.1980.43.3.630. PubMed PMID: 6246217.

37. Huguenard JR, Coulter DA, Prince DA. A fast transient potassium current in thalamic relay neurons: kinetics of activation and inactivation. J Neurophysiol. 1991;66(4):1304–15. Epub 1991/10/01. doi: 10.1152/jn.1991.66.4.1304. PubMed PMID: 1662262.

38. Hodgkin AL, Huxley AF. A quantitative description of membrane current and its application to conduction and excitation in nerve. J Physiol. 1952;117(4):500–44. Epub 1952/08/01. PubMed PMID: 12991237; PubMed Central PMCID: PMCPMC1392413.

39. Hille B. Ion channels of excitable membranes: Sinauer Sunderland, MA; 2001.

40. Jan LY, Jan YN. Voltage-gated potassium channels and the diversity of electrical signalling. J Physiol. 2012;590(11):2591–9. Epub 2012/03/21. doi: 10.1113/jphysiol.2011.224212. PubMed PMID: 22431339; PubMed Central PMCID: PMCPMC3424718.

41. Armstrong CM, Hille B. Voltage-gated ion channels and electrical excitability. Neuron. 1998;20(3):371–80. Epub 1998/04/16. PubMed PMID: 9539115.

42. Pongs O. Voltage-gated potassium channels: from hyperexcitability to excitement. FEBS Lett. 1999;452(1-2):31–5. Epub 1999/06/22. PubMed PMID: 10376673.

43. Staudt N, Molitor A, Somogyi K, Mata J, Curado S, Eulenberg K, et al. Gain-of-function screen for genes that affect Drosophila muscle pattern formation. PLoS Genet. 2005;1(4):e55. Epub 2005/10/29. doi: 10.1371/journal.pgen.0010055. PubMed PMID: 16254604; PubMed Central PMCID: PMCPMC1270011.

44. Bellen HJ, Levis RW, He Y, Carlson JW, Evans-Holm M, Bae E, et al. The Drosophila gene disruption project: progress using transposons with distinctive site specificities. Genetics. 2011;188(3):731–43. Epub 2011/04/26. doi: 10.1534/genetics.111.126995. PubMed PMID: 21515576; PubMed Central PMCID: PMCPMC3176542.

45. Tracey WD, Jr., Wilson RI, Laurent G, Benzer S. painless, a Drosophila gene essential for nociception. Cell. 2003;113(2):261–73. PubMed PMID: 12705873.

46. Kim SE, Coste B, Chadha A, Cook B, Patapoutian A. The role of Drosophila Piezo in mechanical nociception. Nature. 2012;483(7388):209–12. doi: 10.1038/nature10801. PubMed PMID: 22343891; PubMed Central PMCID: PMCPMC3297676.

47. Honjo K, Mauthner SE, Wang Y, Skene JHP, Tracey WD, Jr. Nociceptor-Enriched Genes Required for Normal Thermal Nociception. Cell Rep. 2016;16(2):295–303. doi: 10.1016/j.celrep.2016.06.003. PubMed PMID: 27346357; PubMed Central PMCID: PMCPMC5333372.

48. Robertson RM, Money TG. Temperature and neuronal circuit function: compensation, tuning and tolerance. Curr Opin Neurobiol. 2012;22(4):724–34. doi: 10.1016/j.conb.2012.01.008. PubMed PMID: 22326854.

49. Marder E, Haddad SA, Goeritz ML, Rosenbaum P, Kispersky T. How can motor systems retain performance over a wide temperature range? Lessons from the crustacean stomatogastric nervous system. J Comp Physiol A Neuroethol Sens Neural Behav Physiol. 2015;201(9):851–6. doi: 10.1007/s00359-014-0975-2. PubMed PMID: 25552317; PubMed Central PMCID: PMCPMC4552768.

50. Abrams TW, Pearson KG. Effects of temperature on identified central neurons that control jumping in the grasshopper. J Neurosci. 1982;2(11):1538–53. PubMed PMID: 6292375.

51. Crociani O, Cherubini A, Piccini E, Polvani S, Costa L, Fontana L, et al. erg gene(s) expression during development of the nervous and muscular system of quai. embryos. Mech Dev. 2000;95(1-2):239–43. PubMed PMID: 10906470.

52. Pfeiffer BD, Ngo TT, Hibbard KL, Murphy C, Jenett A, Truman JW, et al. Refinement of tools for targeted gene expression in Drosophila. Genetics. 2010;186(2):735–55. doi: 10.1534/genetics.110.119917. PubMed PMID: 20697123; PubMed Central PMCID: PMCPMC2942869.

53. Szuts D, Bienz M. LexA chimeras reveal the function of Drosophila Fos as a context-dependent transcriptional activator. Proc Natl Acad Sci U S A. 2000;97(10):5351–6. PubMed PMID: 10805795; PubMed Central PMCID: PMCPMC25832.

54. Vacher H, Mohapatra DP, Trimmer JS. Localization and targeting of voltage-dependent ion channels in mammalian central neurons. Physiol Rev. 2008;88(4):1407–47. doi: 10.1152/physrev.00002.2008. PubMed PMID: 18923186; PubMed Central PMCID: PMCPMC2587220.

55. Trimmer JS. Subcellular localization of K+ channels in mammalian brain neurons: remarkable precision in the midst of extraordinary complexity. Neuron. 2015;85(2):238–56. doi: 10.1016/j.neuron.2014.12.042. PubMed PMID: 25611506; PubMed Central PMCID: PMCPMC4303806.

56. Nusser Z. Differential subcellular distribution of ion channels and the diversity of neuronal function. Curr Opin Neurobiol. 2012;22(3):366–71. doi: 10.1016/j.conb.2011.10.006. PubMed PMID: 22033281.

57. Lai HC, Jan LY. The distribution and targeting of neuronal voltage-gated ion channels. Nat Rev Neurosci. 2006;7(7):548–62. Epub 2006/06/23. doi: 10.1038/nrn1938. PubMed PMID: 16791144.

58. Trimmer JS, Rhodes KJ. Localization of voltage-gated ion channels in mammalian brain. Annu Rev Physiol. 2004;66:477–519. Epub 2004/02/24. doi: 10.1146/annurev.physiol.66.032102.113328. PubMed PMID: 14977411.

59. Lerche H, Shah M, Beck H, Noebels J, Johnston D, Vincent A. Ion channels in genetic and acquired forms of epilepsy. J Physiol. 2013;591(4):753–64. Epub 2012/10/24. doi: 10.1113/jphysiol.2012.240606. PubMed PMID: 23090947; PubMed Central PMCID: PMCPMC3591694.

60. Scharfman HE. The neurobiology of epilepsy. Curr Neurol Neurosci Rep. 2007;7(4):348–54. Epub 2007/07/10. PubMed PMID: 17618543; PubMed Central PMCID: PMCPMC2492886.

61. Saras A, Wu VV, Brawer HJ, Tanouye MA. Investigation of Seizure-Susceptibility in a Drosophila melanogaster Model of Human Epilepsy with Optogenetic Stimulation. Genetics. 2017;206(4):1739–46. doi: 10.1534/genetics.116.194779. PubMed PMID: 28630111; PubMed Central PMCID: PMCPMC5560784.

62. Lin WH, Giachello CN, Baines RA. Seizure control through genetic and pharmacological manipulation of Pumilio in Drosophila: a key component of neuronal homeostasis. Dis Model Mech. 2017;10(2):141–50. doi: 10.1242/dmm.027045. PubMed PMID: 28067623; PubMed Central PMCID: PMCPMC5312004.

63. Kuebler D, Zhang H, Ren X, Tanouye MA. Genetic suppression of seizure susceptibility in Drosophila. J Neurophysiol. 2001;86(3):1211–25. doi: 10.1152/jn.2001.86.3.1211. PubMed PMID: 11535671.

64. Lee J, Wu CF. Genetic modifications of seizure susceptibility and expression by altered excitability in Drosophila Na(+) and K(+) channel mutants. J Neurophysiol. 2006;96(5):2465–78. doi: 10.1152/jn.00499.2006. PubMed PMID: 17041230.

65. Marley R, Baines RA. Increased persistent Na+ current contributes to seizure in the slamdance bang-sensitive Drosophila mutant. J Neurophysiol. 2011;106(1):18–29. doi: 10.1152/jn.00808.2010. PubMed PMID: 21451059; PubMed Central PMCID: PMCPMC3129721.

66. Daniels RW, Gelfand MV, Collins CA, DiAntonio A. Visualizing glutamatergic cell bodies and synapses in Drosophila larval and adult CNS. J Comp Neurol. 2008;508(1):131–52. Epub 2008/02/28. doi: 10.1002/cne.21670. PubMed PMID: 18302156.

67. Farooqui T. Review of octopamine in insect nervous systems. Dove Press. 2012;4(1):1–17. doi: https://doi.org/10.2147/OAIP.S20911.

68. Roeder T. Tyramine and octopamine: ruling behavior and metabolism. Annu Rev Entomol. 2005;50:447–77. Epub 2004/09/10. doi: 10.1146/annurev.ento.50.071803.130404. PubMed PMID: 15355245.

69. Ormerod KG, Hadden JK, Deady LD, Mercier AJ, Krans JL. Action of octopamine and tyramine on muscles of Drosophila melanogaster larvae. J Neurophysiol. 2013;110(8):1984–96. doi: 10.1152/jn.00431.2013. PubMed PMID: 23904495.

70. Iliadi KG, Iliadi N, Boulianne GL. Drosophila mutants lacking octopamine exhibit impairment in aversive olfactory associative learning. Eur J Neurosci. 2017;46(5):2080–7. doi: 10.1111/ejn.13654. PubMed PMID: 28715094.

71. Youn H, Kirkhart C, Chia J, Scott K. A subset of octopaminergic neurons that promotes feeding initiation in Drosophila melanogaster. PLoS One. 2018;13(6):e0198362. doi: 10.1371/journal.pone.0198362. PubMed PMID: 29949586; PubMed Central PMCID: PMCPMC6021039.

72. Andrews JC, Fernandez MP, Yu Q, Leary GP, Leung AK, Kavanaugh MP, et al. Octopamine neuromodulation regulates Gr32a-linked aggression and courtship pathways in Drosophila males. PLoS Genet. 2014;10(5):e1004356. doi: 10.1371/journal.pgen.1004356. PubMed PMID: 24852170; PubMed Central PMCID: PMCPMC4031044.

73. Crocker A, Sehgal A. Octopamine regulates sleep in drosophila through protein kinase A-dependent mechanisms. J Neurosci. 2008;28(38):9377–85. doi: 10.1523/JNEUROSCI.3072-08a.2008. PubMed PMID: 18799671; PubMed Central PMCID: PMCPMC2742176.

74. Janigro D, Walker MC. What non-neuronal mechanisms should be studied to understand epileptic seizures? Adv Exp Med Biol. 2014;813:253–64. doi: 10.1007/978-94-017-8914-1_20. PubMed PMID: 25012382; PubMed Central PMCID: PMCPMC4842021.

75. Janigro D, Gasparini S, D’Ambrosio R, McKhann G, 2nd, DiFrancesco D. Reduction of K+ uptake in glia prevents long-term depression maintenance and causes epileptiform activity. J Neurosci. 1997;17(8):2813–24. PubMed PMID: 9092603; PubMed Central PMCID: PMCPMC4066408.

76. Melom JE, Littleton JT. Mutation of a NCKX eliminates glial microdomain calcium oscillations and enhances seizure susceptibility. J Neurosci. 2013;33(3):1169–78. doi: 10.1523/JNEUROSCI.3920-12.2013. PubMed PMID: 23325253; PubMed Central PMCID: PMCPMC3600868.

77. Tanaka K, Watase K, Manabe T, Yamada K, Watanabe M, Takahashi K, et al. Epilepsy and exacerbation of brain injury in mice lacking the glutamate transporter GLT-1. Science. 1997;276(5319):1699–702. PubMed PMID: 9180080.

78. Boison D. Adenosine dysfunction in epilepsy. Glia. 2012;60(8):1234–43. Epub 2012/06/16. doi: 10.1002/glia.22285. PubMed PMID: 22700220; PubMed Central PMCID: PMCPMC3376389.

79. Devinsky O, Vezzani A, Najjar S, De Lanerolle NC, Rogawski MA. Glia and epilepsy: excitability and inflammation. Trends Neurosci. 2013;36(3):174–84. Epub 2013/01/10. doi: 10.1016/j.tins.2012.11.008. PubMed PMID: 23298414.

80. Guizzetti M, Moore NH, Giordano G, Costa LG. Modulation of neuritogenesis by astrocyte muscarinic receptors. J Biol Chem. 2008;283(46):31884–97. Epub 2008/08/30. doi: 10.1074/jbc.M801316200. PubMed PMID: 18755690; PubMed Central PMCID: PMCPMC2581542.

81. Shen JX, Yakel JL. Functional alpha7 nicotinic ACh receptors on astrocytes in rat hippocampal CA1 slices. J Mol Neurosci. 2012;48(1):14–21. Epub 2012/02/22. doi: 10.1007/s12031-012-9719-3. PubMed PMID: 22351110; PubMed Central PMCID: PMCPMC3530828.

82. Araque A, Martin ED, Perea G, Arellano JI, Buno W. Synaptically released acetylcholine evokes Ca2+ elevations in astrocytes in hippocampal slices. J Neurosci. 2002;22(7):2443–50. Epub 2002/03/30. doi: 20026212. PubMed PMID: 11923408.

83. Pabst M, Braganza O, Dannenberg H, Hu W, Pothmann L, Rosen J, et al. Astrocyte Intermediaries of Septal Cholinergic Modulation in the Hippocampus. Neuron. 2016;90(4):853–65. Epub 2016/05/11. doi: 10.1016/j.neuron.2016.04.003. PubMed PMID: 27161528.

84. Haj-Yasein NN, Jensen V, Vindedal GF, Gundersen GA, Klungland A, Ottersen OP, et al. Evidence that compromised K+ spatial buffering contributes to the epileptogenic effect of mutations in the human Kir4.1 gene (KCNJ10). Glia. 2011;59(11):1635–42. doi: 10.1002/glia.21205. PubMed PMID: 21748805.

85. Djukic B, Casper KB, Philpot BD, Chin LS, McCarthy KD. Conditional knock-out of Kir4.1 leads to glial membrane depolarization, inhibition of potassium and glutamate uptake, and enhanced short-term synaptic potentiation. J Neurosci. 2007;27(42):11354–65. doi: 10.1523/JNEUROSCI.0723-07.2007. PubMed PMID: 17942730.

86. Perkins LA, Holderbaum L, Tao R, Hu Y, Sopko R, McCall K, et al. The Transgenic RNAi Project at Harvard Medical School: Resources and Validation. Genetics. 2015;201(3):843–52. Epub 2015/09/01. doi: 10.1534/genetics.115.180208. PubMed PMID: 26320097; PubMed Central PMCID: PMCPMC4649654.

87. Lin DM, Goodman CS. Ectopic and increased expression of Fasciclin II alters motoneuron growth cone guidance. Neuron. 1994;13(3):507–23. PubMed PMID: 7917288.

88. Budnik V, Koh YH, Guan B, Hartmann B, Hough C, Woods D, et al. Regulation of synapse structure and function by the Drosophila tumor suppressor gene dlg. Neuron. 1996;17(4):627–40. PubMed PMID: 8893021; PubMed Central PMCID: PMCPMC4661176.

89. Bier E, Harrison MM, O’Connor-Giles KM, Wildonger J. Advances in Engineering the Fly Genome with the CRISPR-Cas System. Genetics. 2018;208(1):1–18. Epub 2018/01/06. doi: 10.1534/genetics.117.1113. PubMed PMID: 29301946; PubMed Central PMCID: PMCPMC5753851.

90. Bruckner JJ, Zhan H, Gratz SJ, Rao M, Ukken F, Zilberg G, et al. Fife organizes synaptic vesicles and calcium channels for high-probability neurotransmitter release. J Cell Biol. 2017;216(1):231–46. Epub 2016/12/22. doi: 10.1083/jcb.201601098. PubMed PMID: 27998991; PubMed Central PMCID: PMCPMC5223599.

91. Gokcezade J, Sienski G, Duchek P. Efficient CRISPR/Cas9 plasmids for rapid and versatile genome editing in Drosophila. G3 (Bethesda). 2014;4(11):2279–82. doi: 10.1534/g3.114.014126. PubMed PMID: 25236734; PubMed Central PMCID: PMCPMC4232553.

92. Engler C, Kandzia R, Marillonnet S. A one pot, one step, precision cloning method with high throughput capability. PLoS One. 2008;3(11):e3647. doi: 10.1371/journal.pone.0003647. PubMed PMID: 18985154; PubMed Central PMCID: PMCPMC2574415.

93. Zhang X, Koolhaas WH, Schnorrer F. A versatile two-step CRISPR-and RMCE-based strategy for efficient genome engineering in Drosophila. G3 (Bethesda). 2014;4(12):2409–18. doi: 10.1534/g3.114.013979. PubMed PMID: 25324299; PubMed Central PMCID: PMCPMC4267936.

94. Bateman JR, Lee AM, Wu CT. Site-specific transformation of Drosophila via phiC31 integrase-mediated cassette exchange. Genetics. 2006;173(2):769–77. doi: 10.1534/genetics.106.056945. PubMed PMID: 16547094; PubMed Central PMCID: PMCPMC1526508.

95. Chattopadhyay A, Gilstrap AV, Galko MJ. Local and global methods of assessing thermal nociception in Drosophila larvae. J Vis Exp. 2012;(63):e3837. doi: 10.3791/3837. PubMed PMID: 22643884; PubMed Central PMCID: PMCPMC3466948.

